# How prior knowledge prepares perception: Alpha-band oscillations carry perceptual expectations and influence early visual responses

**DOI:** 10.1101/076687

**Authors:** Jason Samaha, Bastien Boutonnet, Bradley R. Postle, Gary Lupyan

## Abstract

Perceptual experience results from a complex interplay of bottom-up input and prior knowledge about the world, yet the extent to which knowledge affects perception, the neural mechanisms underlying these effects, and the stages of processing at which these two sources of information converge, are still unclear. In several experiments we show that language, in the form of verbal cues, both aids recognition of ambiguous “Mooney” images and improves objective visual discrimination performance in a match/non-match task. We then used electroencephalography (EEG) to better understand the mechanisms of this effect. The improved discrimination of images previously labeled was accompanied by a larger occipital-parietal P1 evoked response to the meaningful versus meaningless target stimuli. Time-frequency analysis of the interval between the cue and the target stimulus revealed increases in the power of posterior alpha-band (8-14 Hz) oscillations when the meaning of the stimuli to be compared was trained. The magnitude of the pre-target alpha difference and the P1 amplitude difference were positively correlated across individuals. These results suggest that prior knowledge prepares the brain for upcoming perception via the modulation of alpha-band oscillations, and that this preparatory state influences early (~120 ms) stages of visual processing.

## Introduction

A chief function of visual perception is to “provide a description that is useful to the viewer”^1^, that is, to construct meaning^2,3^. Canonical models of visual perception explain this ability as a feed-forward process, whereby low-level sensory signals are progressively combined into more complex descriptions that are the basis for recognition and categorization^4,5^. There is now considerable evidence, however, suggesting that prior knowledge impacts relatively early stages of perception^6–15^. A dramatic demonstration of how prior knowledge can create meaning from apparently meaningless inputs occurs with two-tone “Mooney” images^16^, which can become recognizable following the presentation of perceptual hints^17,18^.

Although there is general acceptance that knowledge can shape perception, there are fundamental unanswered questions concerning the *type* of knowledge that can exert such effects. Previous demonstrations of Mooney recognition by prior knowledge have used *perceptual* hints, such as pointing out where the meaningful image is located or showing people the completed version of the image^17,19^. Our first question is whether category information cued linguistically— in the absence of any perceptual hints—can have similar effects. Second, it remains unclear whether such effects of knowledge reflect modulation of low-level perception and if so, when during visual processing such modulation occurs. Some have argued that benefits of knowledge on perception reflects late, post-perceptual processes occurring only after processes that could be reasonably called perceptual^20^. In contrast, recent fMRI experiments have observed knowledge-based modulation of stimulus-evoked activity in sensory regions, suggesting an early locus of top-down effects^21–24^. However, the sluggish nature of the BOLD signal makes it difficult to distinguish between knowledge affecting bottom-up processing from later feedback signals to the same regions.

One way that prior knowledge may influence perception is by biasing baseline activity in perceptual circuits, pushing the interpretation of sensory evidence towards that which is expected^25^. Biasing of prestimulus activity according to expectations has been observed both in decision-and motor-related prefrontal and parietal regions ^26–28^ as well as in sensory regions ^21,29,30^. In visual regions, alpha-band oscillations are thought to play an important role in modulating prestimulus activity according to expectations. For example, prior knowledge of the location of an upcoming stimulus changes preparatory alpha activity in visual cortex^31–35^. Likewise, expectations about *when* a visual stimulus will appear are reflected in alpha dynamics ^36–38^. Recently, Mayer and colleagues demonstrated that when the identity of a target letter could be predicted, pre-target alpha power increased over left-lateralized posterior sensors^39^. These findings suggest that alpha-band dynamics are involved in establishing perceptual predictions in anticipation of perception.

Here, we examined whether verbal cues that offered no direct perceptual hints can improve visual recognition of indeterminate two-tone Mooney images (Experiment 1). We then measured whether such verbally ascribed meaning affected an objective visual discrimination task (Experiments 2-3). Finally, we recorded electroencephalography (EEG) during the visual discrimination task (Experiment 4) to better understand the locus at which knowledge influenced perception. Our findings suggest that using language to ascribe meaning to ambiguous images impacts early visual processing by biasing pre-target neural activity in the alpha-band.

## Materials and Method

### Experiment 1

#### Materials

We constructed 71 Mooney images by superimposing familiar images of easily nameable and common artefacts and animals onto patterned background. These superimposed images were then blurred (Gaussian Blur) and then thresholded to a black-and-white bitmap. Materials are available at https://osf.io/stvgy/.

#### Participants

All participants for Experiments 1A-1C were recruited from Amazon Mechanical Turk and were paid $1 (Experiments 1A and 1B), or $0.50 (Experiment 1C) for participating. Demographic information was not collected. All studies were approved by the University of Wisconsin-Madison Institutional Review Board and were conducted in accordance with their policies.

#### Procedure

##### Experiment 1A. Free Naming

We recruited 94 participants (four excluded for non-compliance). Each participant was randomly assigned to view one of 4 subsets of the 71 Mooney images, and to name at the basic-level what they saw in each image. Each image was seen by approximately 24 people. Average accuracies for the 71 images ranged from 0% to 95%.

##### Experiment 1B. Basic Level Cues

From the 71 images used in Experiment 1A we selected the images with accuracy at or below 33% (30 images). We then presented these images to an additional 42 participants (2 excluded for non-compliance. Each participant was shown one of two subsets of the 30 images (15 trials) and asked to choose among 15 basic-level names (e.g., “trumpet”, “leopard”, “table”), which object they thought was present in the image (i.e., a 15-alternative forced choice). Each image received approximately 21 responses.

##### Experiment 1C. Superordinate Cues

Out of the 30 images used in Experiment 1B we selected 15 that had a clear superordinate label (see Fig. 1). Twenty additional participants were presented with each image along with its corresponding superordinate label and were asked to name, at the basic level, the object they saw in their picture by typing their response. For example, given the superordinate cue “musical instrument”, participants were expected to respond with “trumpet” given a Mooney image of a trumpet.

**Fig. 1.**
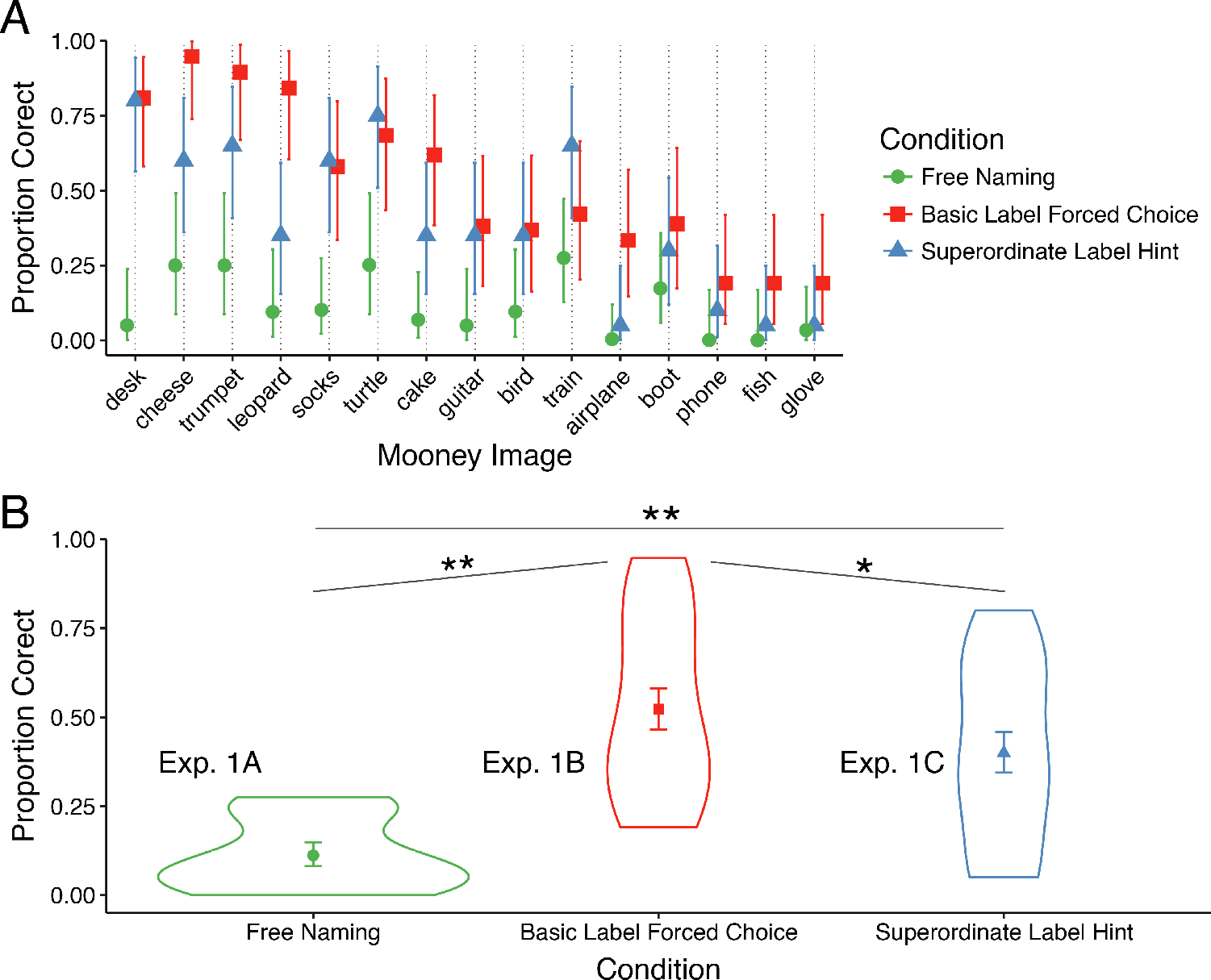
Recognition accuracy from naïve observers (Experiment 1). (**A**) Mean accuracy in the free naming, basic, and superordinate label cue conditions by image. Error bars depict ±1 SEM. (**B**) Violin plot of accuracies averaged over the 15 images used in the three conditions of Experiment 1. Superordinate and basic labels improved recognition accuracy over free naming, and basic labels improved accuracy over superordinate labels. ° Denotes p<.05; ** denotes p<10^−4^

#### Data Coding and Analysis

For free-responses (Experiments 1A, 1C) we considered a response to be correct if it (1) matched the designated image name, e.g., for “lantern”, participants entered “lantern.” (2) was misspelled but identifiable (e.g., “lanturn”), (3) if it was synonymous, e.g., “camping light”, (4) if it contained the target word inside a carrier phrase, e.g., both “socks” and “a pair of socks” was coded as correct. We also coded as correct any “errors” in plurality (e.g., lantern/lanterns) though these were very rare. The responses were first independently coded by three research assistants and any disagreements were discussed until consensus was reached. The effect of condition on accuracy was modeled using logistic regression with a subject and item (Mooney-image category) as random intercepts. The model also included an item-by-condition random slope.

### Experiment 2

#### Materials

From the set of 15 categories used in Experiment 1C, we chose the 10 that had the highest accuracy in the basic-level cue condition (Experiment 1B) and were most benefited by the cues (boot, cake, cheese, desk, guitar, leopard, socks, train, trumpet, turtle). The images subtended approximately 7°×7° of visual angle. Each category (e.g., guitar) was instantiated by four variants: two different image backgrounds and two different positions of the images. These additional images were introduced to tease apart potential detection effects be driven by low-level processing alone.

#### Participants

We recruited 35 college undergraduates to participate in exchange for course credit. Two were eliminated for low accuracy (less than 77%), resulting in 14 participants in the *meaning trained* condition (8 female), and 19 in the *meaning untrained* condition (11 female).

All participants provided written informed consent.

#### Familiarization Procedure

Participants were randomly assigned to a meaning trained or meaning untrained condition. The two conditions differed only in how participants were familiarized with the images. In the meaning trained condition, participants first viewed each Mooney image accompanied by an instruction, e.g., “Please look for CAKE”, twice for each Mooney image (Trials 1-20). Participants then saw all the images again and were asked to type in what they saw in each image, guessing in the case that they could not see anything (Trials 21-30). Finally, participants were shown each image again, asked to type in the label once more and asked to rate on a 1-5 how certain they were that the image portrayed the object they typed. In the meaning untrained condition, participants were familiarized with the images while performing a one-back task, being asked to press the spacebar anytime an image was repeated back-to-back. Repetitions occurred on 20-25% of the trials. In total, participants in the meaning trained and untrained conditions saw each image 4 and 5 times respectively.

#### Same/Different Task

Following familiarization, participants were tested in their ability to visually discriminate pairs of Mooney images. Their task was to indicate whether the two images were physically identical or different in any way (Fig. 2A). Each trial began with a central fixation cross (500 ms), followed by the presentation of one of the Mooney images (the “cue”) approximately 8° of visual angle above, below, to the left or to the right of fixation. After 1500 ms the second image (the “target”) appeared in one of the remaining cardinal positions. The two images remained visible until the participant responded “same” or “different” using the keyboard (hand-response mapping was counterbalanced across participants). Accuracy feedback (a buzz or bleep) sounded following the response, followed by a randomly determined inter-trial interval (blank screen) between 250 and 450 ms. Image pairs were equally divided into three trial-types (Fig 2C): (1) a pair of identical images, (2) a pair of images containing the same object, but in different locations, (3) a pair of images containing different objects at different locations. The backgrounds of the two images on a given trial were always the same. On a given trial, both cue and target objects were either trained or untrained. Participants completed 6 practice trials followed by 360 testing trials and were asked to respond as quickly as possible without compromising accuracy.

**Fig. 2.**
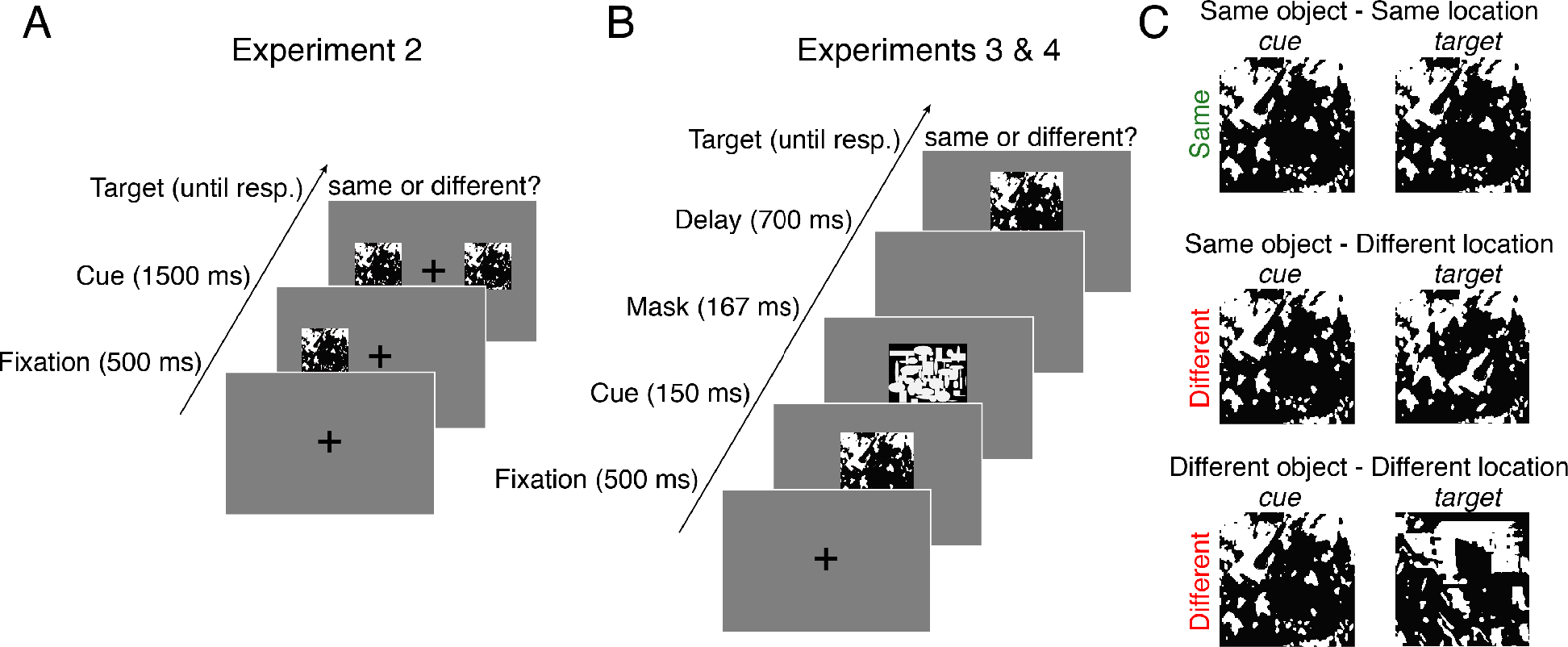
Schematic of the procedure for Experiments 2-4. (**A**) In Experiment 2, participants determined whether two Mooney images were physically identical. (**B**) To increase task difficulty, Experiments 3 and 4 used sequential masked presentation. (**C**) To test for the selectivity of meaning effects, ‘different’ image pairs could differ in object location or object identity. In Experiments 2 and 3, knowledge of the objects was manipulated between participants. In Experiment 4, each participant was exposed to the meanings of a random half of the objects (see Familiarization Procedure).

#### Behavioral Data Analysis

Accuracy was modeled using logistic mixed effects regression with trial-type and meaning-training as fixed effects, subject and item-category random effects with trial-type random slopes. RTs were modeled in the same way, but using linear mixed effects regression. RT analyses excluded responses longer than 5s and those exceeding 3SDs of the subject’s mean.

### Experiment 3

#### Participants

We recruited 32 college undergraduates to participate in exchange for course credit. 16 were assigned to the meaning trained condition (13 female), and the other 16 to the meaning untrained condition (12 female).

#### Familiarization Procedure and Task

The familiarization procedure, task, and materials were identical to Experiment 2 except that the first and second images (approximately 6°×6° of visual angle) were presented briefly and sequentially at the point of fixation, in order to increase difficulty and better test for effects of meaning on task accuracy (see Fig. 2B). On each trial, the initial cue image was presented for 300 ms for the initial 6 practice trials and 150 ms for the 360 subsequent trials. The image was then replaced by a pattern mask for 167 ms followed by a 700 ms blank screen, followed by the target image. Participants’ task, as before, was to indicate whether the cue and target images were identical. The pattern masks were black-and-white bitmaps consisting of randomly intermixed ovals and rectangles (https://osf.io/stvgy/).

#### Behavioral Data Analysis

Exclusion criteria and analysis were the same as in Experiment 2.

### Experiment 4

#### Participants

Nineteen college undergraduates were recruited to participate in exchange for monetary compensation. 3 were excluded from any analysis due to poor EEG recoding quality, resulting in 16 participants (9 female) with usable data. All participants reported normal or corrected visual acuity and color vision and no history of neurological disorders.

#### Familiarization Procedure and Task

The familiarization procedure, task, and materials were nearly identical to that used for Experiment 3, but modified to accommodate a within-subject design. For each participant, 5 of the 10 images were assigned to the meaning trained condition and the remaining to the meaning untrained condition, counterbalanced between subjects. Participants first viewed the 5 Mooney images in the meaning condition together with their names (trials 1-10), with each image seen twice. Participants then viewed the same images again and asked to type in what they saw in each image (trials 11-15). For trials 16-20 participants were again asked to enter labels for the images and prompted after each trial to indicate on a 1-5 scale how certain they were that the image portrayed the object they named. During trials 21-43 participants completed a 1-back task identical to that used in Experiments 2-3 as a way of becoming familiarized with the images assigned to the meaning untrained condition. Participants then completed 360 trials of the same/different task described in Experiment 3.

#### EEG Recording and Preprocessing

EEG was recorded from 60 Ag/AgCl electrodes with electrode positions conforming to the extended 10–20 system. Recordings were made using a forehead reference electrode and an Eximia 60-channel amplifier (Nextim; Helsinki, Finland) with a sampling rate of 1450 Hz. Preprocessing and analysis was conducted in MATLAB (R2014b, The Mathworks, Natick, MA) using custom scripts and the EEGLAB toolbox ^40^. Data were downsampled to 500 Hz offline and were divided into epochs spanning −1500 ms prior to cue onset to +1500 ms after target onset. Epochs with activity exceeding ±75 μV at any electrode were automatically discarded, resulting in an average of 352 (range: 331-360) useable trials per subjects. Independent components responsible for vertical and horizontal eye artifacts were identified from an independent component analysis (using the infomax algorithm with 3 second epochs of 1500 samples each implemented in the EEGLAB function *runica.m*) and subsequently removed. Visually identified channels with poor contact were spherically interpolated (range across subjects: 1-7). After these preprocessing steps, we applied a Laplacian transform to the data using spherical splines ^41^. The Laplacian is a spatial filter (also known as current scalp density) that aids in topographical localization and converts the data into a reference-independent scheme, allowing researchers to more easily compare results across labs; the resulting units are in μV/cm^2^. For recent discussion on the benefits of the surface Laplacian for scalp EEG see ^42,43^.

#### Event-related Potential Analysis

Cleaned epochs were filtered between 0.05 and 25 Hz using a first-order Butterworth filter (MATLAB function *butter.m*). Data were time-locked to target onset, baselined using a subtraction of a 200 ms pre-target window, and sorted according to target meaning condition (trained or untrained). To quantify the effect of meaning on early visual responses, we focused on the amplitude of the visual P1 component. Following prior work in our lab that found larger left-lateralized P1 amplitudes to images preceded by linguistic cues ^44^, we derived separate left and right regions of interest by averaging the signal from occipito-parietal electrodes PO3/4, P3/4, P7/8, P9/10, and O1/2. P1 amplitude was defined as the average of a 30 ms window, centered on the P1 peak as identified from the grand average ERP (see Fig. 4A).

**Fig. 3.**
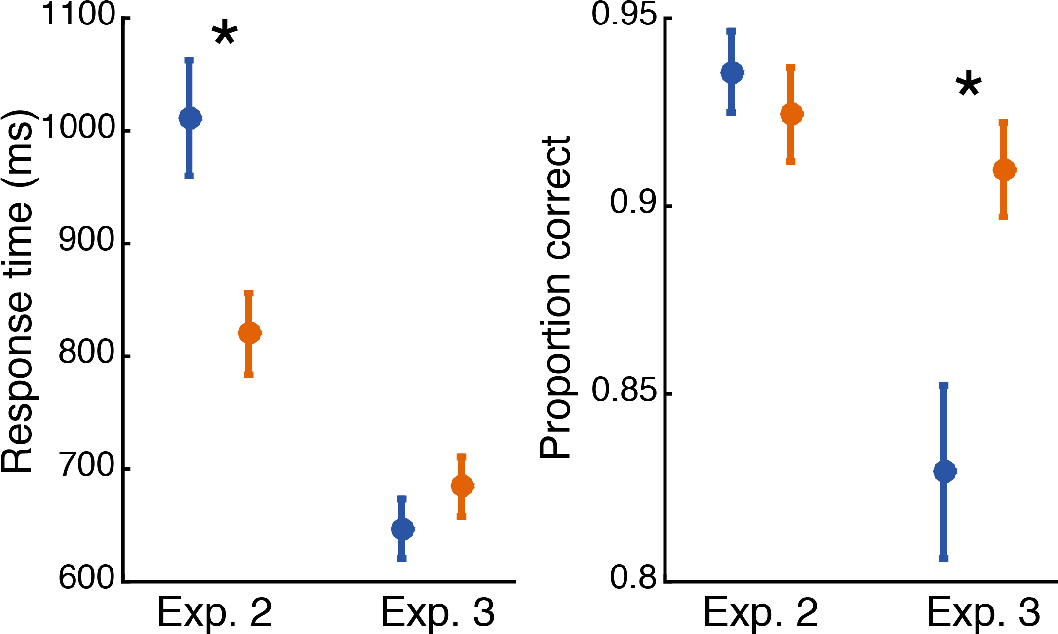
Response time (left panel) and accuracy (right panel) for Experiments 2 and 3. Meaning training significantly decreased response time in Experiment 2 (when both images were presented simultaneously and remained visible until response), and significantly improved accuracy in Experiment 3 (when images were presented briefly and sequentially). Error bars show ±1 SEM; asterisks indicated two-tailed significance at p<.05.

**Fig. 4.**
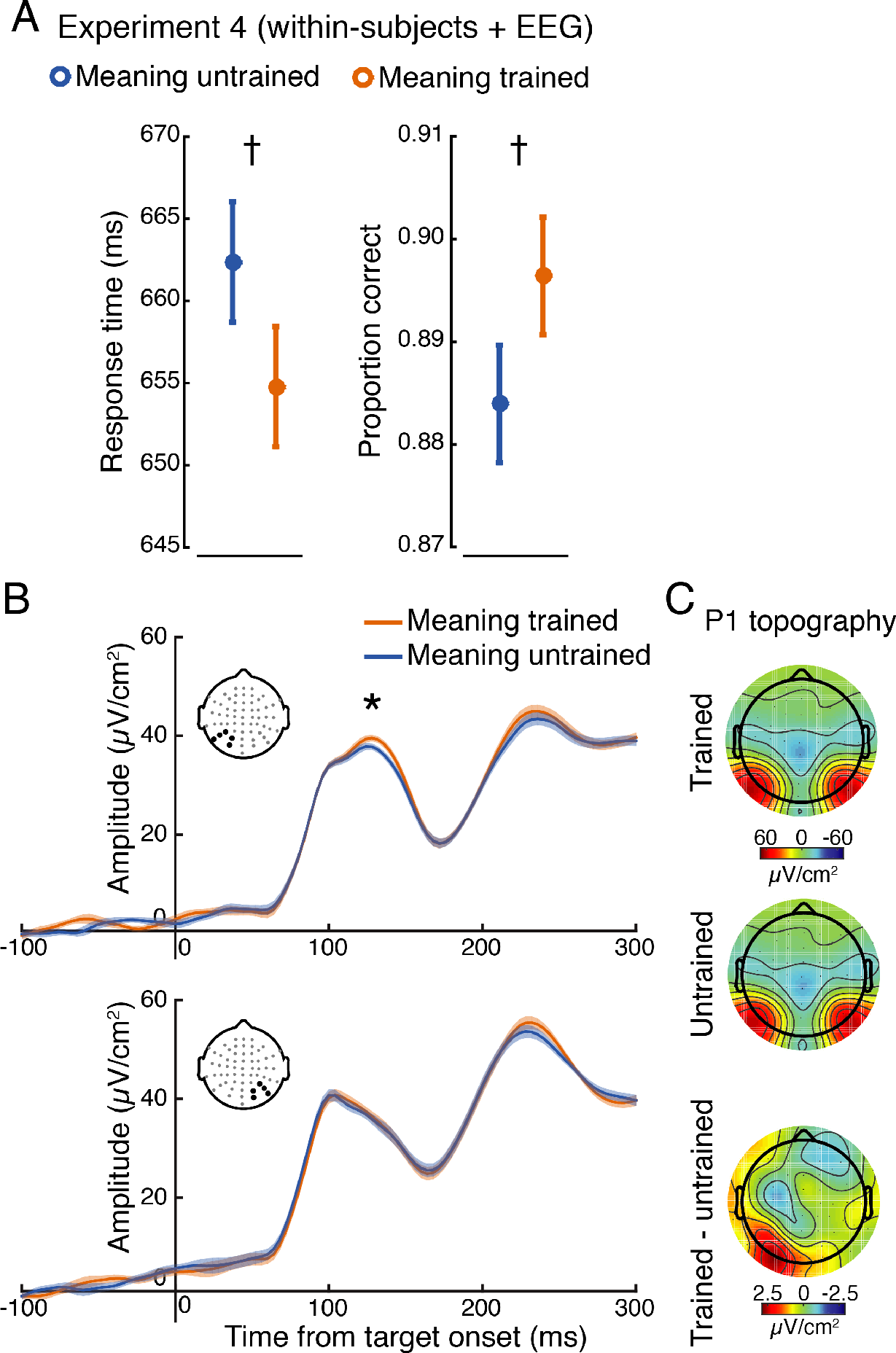
Behavior and electrophysiology results from Experiment 4. (**A**) Response time (left panel), accuracy (right panel) showed trending improvements for images previously made meaningful. Note that a model including both accuracy and RTs revealed a significant effect of meaning training on accuracy (see Results). (**B**) Analysis of the P1 event-related potential revealed a significant main effect, indicating larger amplitude responses to meaning trained targets. This main effect was largely driven by significant differences at left posterior electrodes (upper panel; signal averaged over electrodes denoted with black dots), but not right (lower panel), although the interaction did not reach significance. (**C**) Topography of the P1 for both conditions and their difference. Error bars and shaded bands represent ±1 within-subjects SEM ^84^; asterisks indicated two-tailed significance at p<.05; daggers represent two-tailed trends at p<.08.

This same procedure was used to analyze P1 amplitudes in response to the cue stimulus, with the exception that baseline subtraction was performed using the 200 ms prior to cue onset. Lastly, in order to relate P1 amplitude and latencies to behavior, we used a single-trial analysis. As in prior work ^44^, single-trial peaks were determined from each electrode cluster (left and right regions of interest) by extracting the largest local voltage maxima between 70 to 150 ms post-stimulus (using the MATLAB function *findpeaks*). Any trial without a detectable local maximum (on average ~1%) was excluded from analysis.

#### Time-Frequency Analysis

Time-frequency decomposition was performed by convolving single trial unfiltered data with a family of Morelet wavelets, spanning 3–50 Hz, in 1.6-Hz steps, with wavelet cycles increasing linearly between 3 and 10 cycles as a function of frequency. Power was extracted from the resulting complex time series by squaring the absolute value of the time series. To adjust for power-law scaling, time-frequency power was converted into percent signal change relative to a common condition pre-cue baseline of −400 to −100 ms. To identify time-frequency-electrode features of interest for later analysis in a data-driven way while avoiding circular inference, we first averaged together all data from all conditions and all electrodes. This revealed a prominent (~65% signal change from baseline) task-related increase in alpha-band power (8-14 Hz) during the 500 ms preceding target onset, with a clear posterior scalp distribution (see Fig. 5A), in-line with the topography of alpha observed in many other experiments ^45,46^. Based on this, we focused subsequent analysis on 8-14 Hz power across the pre-target window −500 to 0 ms using the same left/right posterior electrode clusters as in the ERP analysis.

**Fig. 5.**
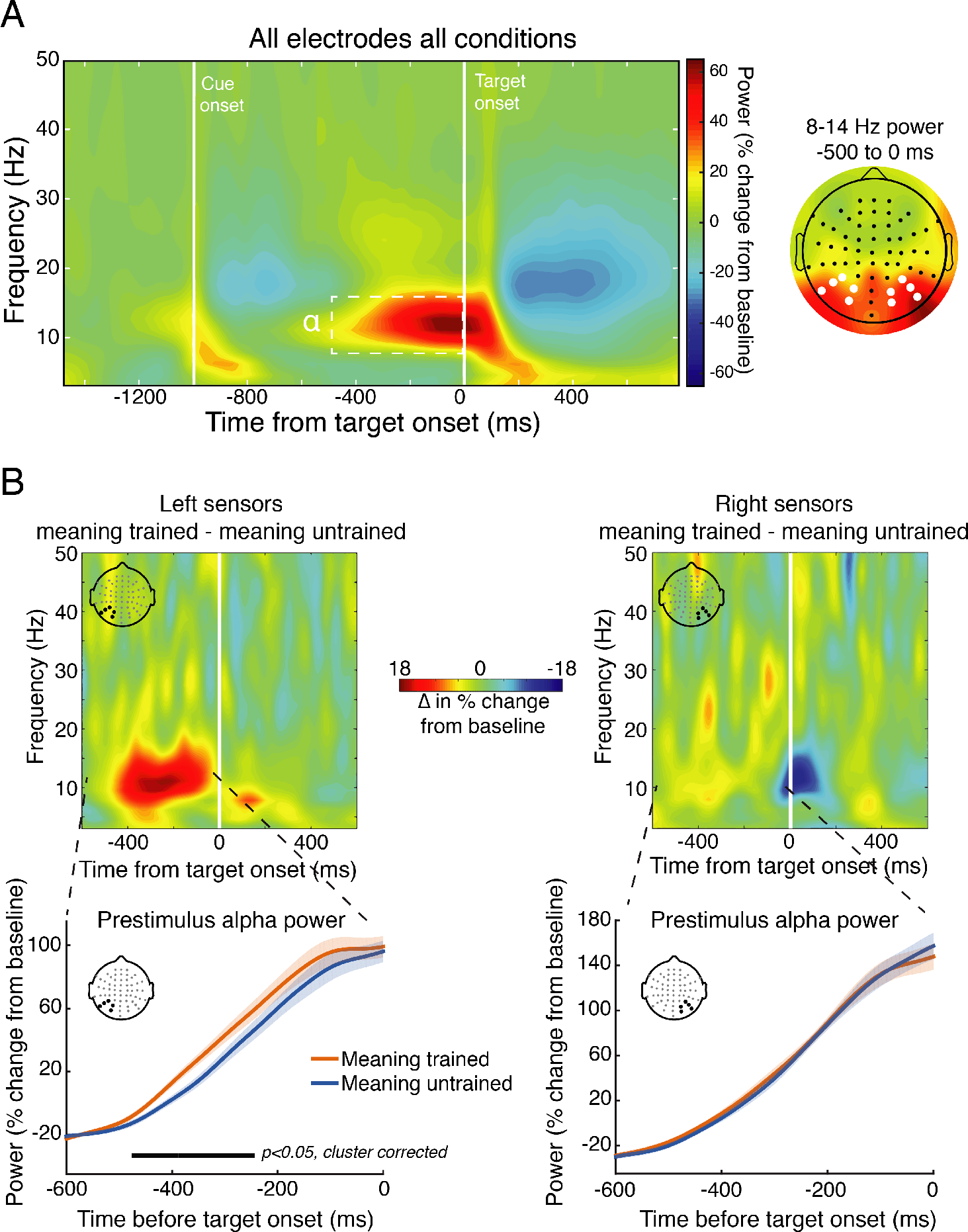
Time-frequency analysis of alpha-band power during the cue-target interval (Experiment 4). (**A**) To identify time-frequency-electrode regions of interest while avoiding circular inference, we averaged time-frequency power across all electrodes and conditions. This revealed a prominent increase (~ 65% from baseline) in pre-target (−500 to 0 ms) power in the alpha range (8-14 Hz) that had a posterior topography (right panel; left and right electrode clusters of interest denoted with white dots) associated with simply performing the task. We then focused on how meaning training impacted this signal in subsequent analyses. (**B**) Time-frequency power plots showing the difference (meaning-trained – meaning-untrained) for left (left panel) and right (right panel) electrodes of interest (derived from panel A) reveal greater alpha power just prior to target onset on meaning-trained trials. The lower panels depict the time-course of the pre-target alpha signal for meaning-trained and untrained trials, revealing a significant temporal cluster of increased alpha power approximately 480 to 250 ms prior to target onset over left, but not right electrode clusters. Shaded regions represent ±1 within-subjects SEM.

#### Statistical Analysis

We conducted two analyses of pre-target alpha power. To examine the effect of meaning training on the time course of pre-target alpha power (see Fig. 5B), we analyzed left and right electrode groups separately with a non-parametric permutation test and cluster correction to deal with multiple comparisons across time points ^47^. This was accomplished by randomly shuffling the association between condition labels (meaning trained or untrained) and alpha power 10,000 times. On every iteration, a *t*-statistic comparing alpha power between meaning trained and meaning untrained conditions was computed for each time sample. The largest number of contiguous significant samples was saved, forming a distribution of *t*-statistics under the null hypothesis that meaning training had no effect, as well as a distribution of cluster sizes expected under the null. The *t*-statistic associated with the true data mapping was compared, at each time point, against this null distribution and only cluster sizes exceeding the 95% percentile of the null cluster distribution was considered statistically different. α was set at 0.05 for all comparisons. In the second analysis which additionally tested for an interaction between hemispheres, we averaged alpha power across the pre-target window −500 to 0 ms and fit a linear mixed-effects model using meaning condition (trained vs. untrained), electrode cluster (left vs. right hemisphere), and their interaction to predict alpha power, with random slopes for meaning condition and hemisphere by subject (this model is equivalent to a 2-by-2 repeated-measures ANOVA).

To predict trial-averaged P1 amplitudes we used a linear mixed-effects model predicting P1 amplitude from meaning (trained vs. untrained), electrode cluster (left vs. right hemisphere), and their interaction, with random slopes for meaning condition and hemisphere by subject. Simple effects were then tested using paired *t*-tests to compare P1 amplitudes and pre-target alpha power between meaning conditions separately for each electrode group. We examined simple effects on the basis of two recent reports examining the influence of linguistic ^44^ and perceptual cues ^39^ on P1 amplitudes. Both of these experiments found left-lateralized P1 enhancements to cued images. We therefore anticipated significant differences over left, but not right sensors, and report simple effects in addition to main effects and interactions. Regarding the single-trial P1 analysis (see *Event-related Potential Analysis* above), we used linear mixed-effects models with subject and item random effects to examine the relationship between single-trial P1 peak amplitudes and latencies to the accuracy and latency of behavioral responses. See https://osf.io/stvgy/ for full model syntax. Where correlations are reported, we used Spearman rank coefficients to test for monotonic relationships while mitigating the influence of potential outliers. We additionally conducted a non-parametric bootstrap analysis (20000 bootstrap samples) to form 95% confidence intervals around across-subject correlation coefficients and to verify the significance of any correlation using an additional non-parametric statistic.

## Results

### Experiment 1

Mean accuracy for the 15 images used in all versions of Experiment 1 is displayed in Fig. 1A; the means for the three naming conditions (Experiments 1A, 1B, and 1C, respectively) are shown in Fig. 1B. (For accuracy results of the remaining Mooney images, see https://osf.io/stvgy/). Baseline recognition performance (free-naming; Experiment 1A) was 24.4% for the full set of 71 images and 11% (95% CI=[.08, .15]) for the set of 15 used in all three versions of Experiment 1. Providing participants with a list of 15 possibilities (Experiment 1B) increased recognition from 11% to 52% (95% CI=[.47, .58]). A logistic regression analysis revealed this to be a highly significant difference (b=2.74, 95% CI=[1.94, 3.54], z=6.7, p<10^−4^). Part of this increase in Experiment 1b is likely due to the difference in the response formats between Experiments 1A (free response) and 1B (multiple choice with 15 simultaneously presented options). Experiment 1C used the free-response format of Experiment 1A, but provided participants a non-perceptual hint in the form of a superordinate label (e.g., “animal”, “musical instrument”). This simple hint yielded recognition performance of 40% (95% CI=[.34, .46]), a nearly 4-fold increase compared to baseline free-response (b=1.92, 95% CI=[1.22, 2.61], z=5.39, p<10^−4^). For example, knowing that there is a piece of furniture in the image produced a 16-fold increase in accuracy in recognizing it as a desk (an impressive result even allowing for guessing). Providing basic-level alternatives (Experiment 1B) yielded significantly greater performance than providing superordinate-hints (b=.73, 95% CI=[.13, 1.32], z=2.40, p=.02), although this difference is difficult to interpret owing to a difference in the response format between the two tasks. The main conclusion from Experiment 1 is that recognition of two-tone images can be drastically improved by verbal hints that provide no spatial or other perceptual information regarding the identity of the image.

### Experiment 2

Results are shown in Fig. 3. Overall accuracy was high—93.3% (93.8% on trials showing two different images and 92.2% on trials showing identical image pairs). The accuracy for the meaning-trained participants (M=92.6%) was not significantly different from the participants not trained on meanings (M=93.4%; b=−.18, 95% CI=[−.75, .40], z=−0.62, p=.54). Given the ease of the perceptual discrimination task and participants had unlimited time to inspect the two images, an absence of an accuracy effect is not surprising. Participants in the meaning-trained condition, however, had significantly shorter RTs than those who were not exposed to image meanings: RT_meaning trained_=822 ms; RT_meaning untrained_=1017ms (b=-194, 95% CI=[−326, −61], t=−2.86, p<.01; see Fig. 3). There was a marginal trial-type by meaning interaction (b=71, 95% CI=[−1, 144], t=1.93, p=.06). Meaning was most beneficial in detecting that two images were exactly identical, (b=−262, 95% CI=[−445, −79], t=−2.80, p<.01). There remained a significant benefit of meaning in detecting difference in images with the same object in a different location, (b=−201, 95% CI=[−353, −49], t=2.60, p=.01) and a smaller difference when two images had different objects and object locations, (b=−121, 95% CI=[−219, −23], t=−2.41, p=.02).

### Experiment 3

The brief presentation of the cue-image in Experiment 3 (Fig. 2B) had an expected detrimental effect on accuracy, which was now 87.2% (90.0% on different trials and 81.7% on same trials), significantly lower than accuracy of Experiment 2 (b=−.75, 95% CI=[−1.12, −0.38], z=−3.91, p<10^−4^). Participants’ responses were significantly faster (M=664 ms) than in Experiment 2 (b=− .264, 95% CI=[−349, −179], t=−6.1, p<10^−4^). This may seem odd given the greater difficulty of the procedure, but unlike Experiment 2 in which participants could improve their accuracy by spending additional time examining the two images, in the present study performance was limited by how well participants could extract and retain information about the brief cue image.

Exposing participants to the image meanings significantly improved accuracy: M_meaning_ trained=91.3%; M_meaning untrained_=83.1% (b=.71, 95% CI=[.26, 1.17], z=3.06, p<.01; Fig. 3). The meaning advantage interacted significantly with trial type (b=.33, 95% CI = [.10, .55], z=2.87, p<.01). The increase in accuracy following meaning training was again largest for the identical-image trials (b=1.12, 95% CI=[.60, 1.64], z=4.22, p<10^−4^). It was smaller when the two images showed the same object in different locations (b=.57, 95% CI=[.08, 1.07] z=2.30, p=.02), and marginally so when the two images showed different objects in different locations (b=.70, 95% CI=[−.09, 1.49], z=1.73, p=.08).

Meaningfulness did not significantly affect RTs, which were slightly longer for meaning-trained participants (M=685 ms) than meaning-untrained participants (M=648 ms). Although this was far from reliable, b=36, 95% CI =[−328, 61], t=.80, p=.43, we sought to check that the accuracy advantage reported above still obtained when RTs were taken into account. We therefore included RT (on both correct and incorrect trials) as an additional fixed predictor in the logistic regression. RT was strongly related to accuracy: faster responses corresponded to greater accuracy, b=−.0010, 95% CI=[ −0.0012, −0.0009], z=−13.3, p<10^−4^, i.e., there was no evidence of a speed-accuracy tradeoff. Meaning-training remained a significant predictor of accuracy when RTs were included in the logistic regression as a fixed effect, b=.77, 95% CI=[.28, 1.26], z=3.08, p<.01.

### Experiment 4

#### Behavior

Overall accuracy was 89.0% (92.8% on different trials and 81.3% on same trials). Participants were marginally more accurate when discriminating images previously made meaningful compared to images whose meaning was untrained: M_meaning-trained_=89.8%; M_meaning-untrained_=88.2% (b=.21, 95% CI=[−0.0001, .42], z=1.96, p=.05; Fig. 4A). The meaning-by-trial-type interaction for accuracy was not significant, p>.8. Overall RT was, at 641 ms— comparable to Experiment 3—and was marginally shorter when discriminating images that were previously rendered meaningful: M_meanin-trained_=656 ms; M_meaning-untrained_=665 ms, (b=−9.7, 95% CI=[−21, 1.1], t=−1.76, p=.08; Fig. 4A). The meaning-by-trial-type interaction for RTs was not significant, p>.90. As evident from Fig 4A, the effect of meaning-training was split between accuracy and RTs. We therefore repeated the accuracy analysis including RT (for both correct and incorrect trials) as an added predictor. As in Experiment 3, RTs were negatively correlated with accuracy, b=−.0012, 95% CI=[−.0016, −.0010], t=−9.17, p<.10^−4^. With RTs included in the model, meaning-training was associated with greater accuracy, b=.22, 95% CI=[.012, .43], t=2.08, p=.04.

Combining experiments 3 and 4 revealed a significant effect of meaning-training on accuracy b=.57, 95% CI=[0.24, 0.90], t=3.35, p<10^−3^, and a significant meaning-training by experiment interaction b=−.64, 95% CI=[−1.20, −.08], t=−2.21, p=.03, suggesting that the effect of meaning-training on accuracy was larger in experiment 3 compared to experiment 4. Including RT in the model did not appreciably change these results. We speculate that the reduced effect in the present experiment is due to the within-subject manipulation of meaningfulness.

#### P1 amplitude analysis

As shown in Fig. 4B, trial-averaged P1 amplitude was significantly larger when viewing targets whose meaning was trained, as compared to those whose meaning was untrained (b=−1.7, 95% CI=[−3.29, −0.13], t=−2.16, p=.037). Although there was no significant interaction with hemisphere (p=.22), analysis of simple effects using paired *t*-tests revealed that meaning increased P1 amplitudes at the left hemisphere electrode cluster (t(15)=2.59, 95% CI=[0.30, 3.12], p=.02), but not at right (t(15)=.35, 95% CI=[−1.68, 2.36], p=.72). These same analyses were repeated for cue-evoked P1 amplitudes. No main effect or interaction was observed (both p-values>.70), suggesting that the effect of meaning on P1 amplitudes was specific to the target-evoked response.

#### Pre-target Alpha-band Power

The linear mixed-effects model of alpha power (averaged over the 500 ms prior to target onset) revealed a significant effect of meaning (b=−9.85, 95% CI=[−18.42, −1.29], t=−2.3, p=.03), indicating greater pre-target alpha power on meaning trained trials, and a significant interaction between hemisphere and meaning (b=8.31, 95% CI=[ 2.27, 14.36], t=2.75, p=.01). Paired *t*-tests revealed that meaning increased pre-target alpha power in the left (t(15)=2.21, 95% CI=[0.33, 19.38], p=.04), but not right (t(15)=0.35, 95% CI=[−7.78, 10.86], p=.72) hemisphere. Analysis of the time course of pre-target alpha power revealed a significant cluster-corrected increase in power on meaning-trained trials from approximately −480 to −250 ms prior to target onset. Significant clusters were observed over left occipito-parietal sensors, but not right (see Fig. 5B). Note that this pre-target difference is unlikely to be accounted for by temporal smoothing of post-target differences as there were clearly no post-target differences (Fig. 5B).

#### Alpha Power and P1 Correlation

We next assessed the relationship between the meaning effect on pre-target alpha power and on P1 amplitudes across participants by correlating alpha modulations (averaged over the pre-target window) with P1 modulations, for both right and left electrode groups. This analysis revealed a significant positive correlation (rho=0.52, p=.04, bootstrap 95% CI=[0.08, 0.82]) over left electrodes, indicating that individuals who showed a greater increase in pre-target alpha from meaning training also had a larger effect of meaning on P1 amplitudes (see Fig. 6A). This relationship was not significant over right hemisphere electrodes (rho = −0.21, p=.42, bootstrap 95% CI=[−0.71, 0.41]; Fig. 6B). These two correlations were significantly different (p=.04) and the 95% CI of the difference between bootstrap distributions only slightly overlapped with zero (CI=[1.39, −0.04]), suggesting that these interactions may be specific to the left hemisphere.

**Fig. 6.**
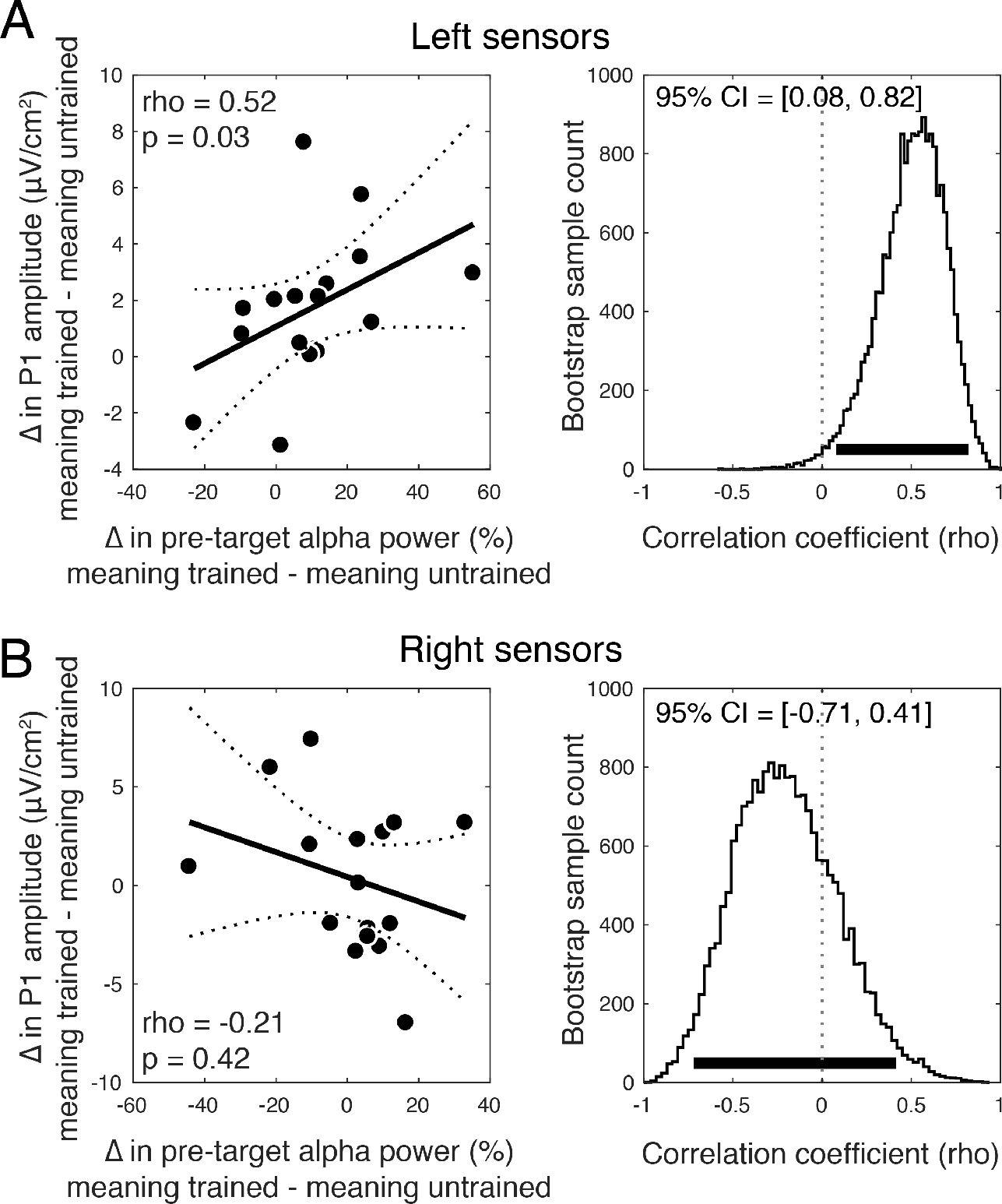
The magnitude of the meaning training effect on pre-target alpha power predicts the magnitude of the meaning effect on P1 amplitude across participants. (**A**) A significant positive correlation over left hemisphere sensors indicates that individuals who showed a greater increase in pre-target alpha power on meaning-trained trials also showed a greater increase in P1 amplitude. Dashed lines denote the 95% CI on the linear fit and the right panel shows bootstrap distributions of the correlation coefficient with 95% CI denoted with a thick black line. (**B**) Same correlation but for right hemisphere electrodes, which was non-significant and with a bootstrap distribution substantially overlying zero.

#### Single-trial P1 Analysis

Finally, we used linear mixed-effects models with subject and item random effects to examine the relationship between single-trial P1 peak amplitudes and latencies and the accuracy and latency of participants’ same/different responses. We focused on P1 peaks from the cluster of left electrodes because these sensors were driving the significant P1 maineffect at the trial-averaged model (see above), as well as the significant alpha power interaction. A focus on left posterior electrodes was also warranted by work in our lab that found P1 modulation by linguistic cues occurring over left occipito-parietal sensors^44^.

Per-trial amplitudes were numerically greater for meaningful trials (*M*=64.43 μV/cm^2^) than meaningless trials (M=63.70 μV/cm^2^), but not significantly so, b=.83, 95% CI = [−.82, 2.50], t=1.02, p=.31. There was a significant interaction between behavioral RTs and meaningfulness in predicting the P1 amplitude, b= −0.008, 95% CI=[−0.014 −0.001], t=−2.48, p=.01, such that on meaningful trials, larger P1s were associated with faster behavioral responses (controlling for accuracy), b=−0.008, 95% CI=[−.013, −.002], t=2.68, p<.01. On meaningless trials, no such relationship was observed, b= 0.0006, 95% CI=[−0.005, 0.006], t=.24, p>.40. There was no relationship between P1 peak amplitude and accuracy either for meaningful or meaningless trials, t’s<1.

Intriguingly, P1 latencies were slightly, but significantly *delayed* on meaningful (M=114.8 ms) compared to meaningless trials (M=113.4 ms), b=1.50, 95% CI=[.50, 2.50], t=2.96, p<.01. A later P1 seems suboptimal, yet within subjects, later P1s were associated with shorter RTs, b=−0.003, 95% CI = [−.0054, −.0005], t=2.33, p=.02 (no interaction with meaningfulness was observed, t=−.32). As with P1 amplitude, P1 latencies were uncorrelated with accuracy for either meaningful or meaningless trials t’s<1.

Unlike left-hemisphere electrodes, per-trial P1 amplitudes over the right electrodes were numerically smaller for meaningful trials (M=71.34 μV/cm2) than meaningless trials (M=71.97 μV/cm2), but not significantly so, b=.83, 95% CI = [−.82, 2.50], t<1. There was no significant interaction between behavioral RTs and meaningfulness in predicting the P1 amplitude, t<1 Similar to the left hemisphere electrodes, however, P1 latencies were longer on meaningful trials (M=110.8 ms) compared to meaningless trials (109.3 ms), b=1.52, t=2.93, p=.004, 95% CI= [.50, 2.52]. Unlike the left-hemisphere electrodes, however, these longer P1 latencies on meaningful trials were not significantly associated with behavioral RTs, b=.002, t=1.23, p=.22. Full analyses can be found at https://osf.io/stvgy/.

#### Control Analyses

To determine whether participants’ improved performance for the meaning trained images could be explained by learning *where* the object was located and looking to those locations we analyzed electrooculograms (EOGs, prior to ocular correction from ICA) recorded from bipolar electrodes placed on the lateral canthus and lower eyelid of each participant’s right eye during the EEG recording. If participants more frequently engaged in eye movement during the cue-target interval of meaning-trained trials we would expect, on average, larger amplitude EOG signals following the cue. However, EOG amplitudes, time-locked to the onset of the cue, did not reliably distinguish between meaning-trained and meaning-untrained trials in the way that alpha power during this same interval did (all p-values>.65, time-cluster corrected). EOG amplitudes on meaning-trained trials also did not reliably differ when trials were sorted by the location of the object in the cue image: whether it was on the left or right side, on the top or bottom, or lateral or vertical relative to center (all p-values>.43, time-cluster corrected). Across the whole cue-target interval, no contrast survived the same cluster correction procedure applied to the alpha time-course analysis, suggesting that eye movements are unlikely to explain our EEG findings.

To investigate the possibility that participants covertly attended to the location of the object in the cue image, we tested for well-known effects of spatial attention on alpha lateralization. Numerous studies have demonstrated alpha power desynchronization at posterior electrodes contralateral to the attended location ^31,32,34^. Thus, if subjects were maintaining covert attention, for example, to the left side of the image following a cue with a left object, then alpha power may decrease over right sensors relative to when a cue has an object on the right, and vice versa. Contrary to this prediction, we observed no modulation of alpha power at either left (all p-values>.94, time-cluster corrected) or right electrode clusters (all p-values>.35, time-cluster corrected) by the object location within the Mooney image. This suggests that spatial attention is not the source of the effects of meaning training.

To ensure that the P1 effect and the across-subject correlation between alpha power and P1 were not dependent on filter choices applied during preprocessing, we re-conducted both analyses using unfiltered data. Regarding the P1, we again observed a main effect of meaning training on P1 amplitudes (b=−1.8, 95% CI=[−3.42, −0.20], t=−2.25, p=.030), indicating larger P1’s following meaning-trained targets and no main effect of hemisphere, or interaction (t’s<0.9). Paired *t*-tests confirmed that P1 amplitudes were larger for meaning-trained targets at left hemisphere electrodes (t(15)=2.51, 95% CI=[0.27, 3.35], p=.02), but not at right (t(15)=.76, 95% CI=[−1.26, 2.68], p=.45). Regarding the correlation between pre-target alpha power and P1 modulations, we again found a significant across-subject correlation at left hemisphere electrodes (rho=0.61, p=0.01, bootstrap 95% CI=[0.21, 0.85]), but not at right (rho=−0.26, p=.32, bootstrap 95% CI=[−0.74, 0.37]). These correlations were significantly different from one another as the 95% CI of the difference between left and right electrode bootstrap distributions did not contain zero (CI=[1.43, 0.051]).

## Discussion

To better understand when and how prior knowledge influences perception we first examined how non-perceptual cues influence recognition of initially meaningless Mooney images. These verbal cues resulted in substantial recognition improvements. For example, being told that an image contained a piece of furniture produced a 16-fold increase in recognizing a desk (Fig.1). We next examined whether ascribing meaning to the ambiguous images improved not just people’s ability to *recognize* the denoted object, but to perform a basic perceptual task: distinguishing whether two images were physically identical. Indeed, ascribing meaning to the images through verbal cues improved people’s ability to determine whether two simultaneously or sequentially presented images were the same or not (Fig. 3 and 4). The behavioral advantage might still be thought to reflect an effect of meaningfulness on some relatively late process were it not for the electrophysiological results showing that ascribing meaning led to increase in the amplitude of P1 responses to the target (Fig. 4B)^cf. 48^. The P1 enhancement was preceded by an increase in alpha amplitude during the cue-target interval when the cue was meaningful (Fig. 5). The effect of meaning training on pre-target alpha power and target-evoked P1 amplitude were positively correlated across participants, such that individuals who showed larger increases in pre-target alpha power as a result of meaning training, also showed larger increases in P1 amplitude (Fig. 6). Combined, our results contradict claims that knowledge affects perception only at a very late stage ^49,20,50^ and provide general support for predictive processing accounts of perception, positing that knowledge may feedback to modulate lower levels of perceptual processing.^3,25,51^.

In Experiment 2, when meaning training was manipulated between subjects and participants could compare both images with unlimited time we observed effects of meaning on RTs but not accuracy. When the visual discrimination was made difficult via masking and brief presentation times (Experiments 3 and 4), effects on accuracy were more pronounced. This was true for both between-and within-subject versions of the manipulation (Experiments 3 and 4, respectively). However, there were notable differences between behavioral performance in Experiments 3 and 4. The meaning effect on accuracy in Experiment 4 was reduced compared to Experiment 3 and a trending response time effect emerged in Experiment 4. Additionally, there was an interaction with trial type and meaning predicting accuracy in Experiment 3, but not Experiment 4. These differences are possibly due, in part, to the change from between-subjects to within-subjects in Experiment 4 which could have resulted in some of the meaning untrained images being recognized due to exposure to both conditions. That is, the effectiveness of the meaning manipulation may have been reduced as a result of all the subjects in this experiment knowing that the stimuli contained meaningful objects.

These behavior results are novel in two respects. First, it marks the first demonstrations, to our knowledge, of cuing recognition of Mooney-style images using solely linguistic cues, as opposed to the more common method of simply revealing the original image ^17,18,52^. Second, the results of our same/different discrimination task reveal that linguistic cues enhance not only the ability to recognize the images, as in prior work, but also putatively lower-level processes subserving visual discrimination.

The P1 ERP component is associated with relatively early regions in the visual hierarchy (most likely ventral peri-striate regions within Brodmann’s Area 18 ^53–56^) but is has been shown to be sensitive to top-down manipulations such as spatial cueing ^57,58^, object based attention ^59^, object recognition ^60,61^, and recently, trial-by-trial linguistic cueing ^44^. Our finding that averaged P1 amplitudes were increased following meaning training is thus most parsimoniously explained as prior knowledge having an early locus in its effects on visual discrimination (although the failure to find this effect in the single-trial EEG suggests some caution in its interpretation). This result is consistent with prior fMRI findings implicating sectors of early visual cortex in the recognition of Mooney images ^17,52^ but extends these results by demonstrating that the timing of Mooney recognition is consistent with the modulation of early, feedforward visual processing. Interestingly, the effect of meaning on P1 amplitude was present only in response to the target stimulus, and not the cue. This suggests that, in our task, prior knowledge impacted early visual responses in a dynamic manner, such that experience with the verbal cues facilitated the ability to form expectations for a subsequent “target” image. We speculate that this early target-related enhancement may be accomplished by the temporary activation of the cued perceptual features (reflected in sustained alpha power) rather than by an immediate interaction with long-term memory representations of the meaning-trained features, which would be expected to lead to enhancements of both cue and target P1. Another possibility is that long-term memory representations are brought to bear on the meaning-trained “cue” images, but these affect later perceptual and post-perceptual processes.

Our findings are also in line with two recent magnetoencephalography (MEG) studies reporting early effects of prior experience on subjective visibility ratings ^39,62^. In those studies, however, prior experience is difficult to disentangle from perceptual repetition. For example, Aru and colleagues ^62^ compared MEG responses to images that had previously been studied against images that were completely novel, leaving open mere exposure as a potential source of differences. In our task, by contrast, participants were familiarized with both meaning trained and meaning untrained images but only the identity of the Mooney image was revealed in the meaning training condition, thereby isolating effects of recognition. Our design further rules out the possibility that stimulus factors (e.g., salience) could explain our effects, since the choice of which stimuli were trained was randomized across subjects. One possible alternative by which meaning training may have had its effect is through spatial attention. For example, it is conceivable that on learning that a given image has a boot on the left side, participants subsequently were more effective in attending to the more informative side of the image. If true, such an explanation would not detract from the behavioral benefit we observed, but would mean that the effects of knowledge were limited to spatial attentional gain. Subsequent analyses suggest this is not the case (see *Control Analyses*).

It is noteworthy that, as in the present results, the two MEG studies mentioned above, as well as related work from our lab employing linguistic cues^44^, have all found early effects over left-lateralized occipito-parietal sensors, suggesting that the effects of linguistically aided perception may be more pronounced in the left hemisphere, perhaps owing to the predominantly left lateralization of lexical processing^63^.

Mounting neurophysiological evidence has linked low-frequency oscillations in the alpha and beta bands to top-down processing ^64–67^. Recent work has demonstrated that perceptual expectations modulate alpha-band activity prior to the onset of a target stimulus, biasing baseline activity towards the interpretation of the expected stimulus ^28,39^. We provide further support for this hypothesis by showing that posterior alpha power increases when participants have prior knowledge of the meaning of the cue image, which was to be used as a comparison template for the subsequent target. Further, pre-target alpha modulation was found to predict the effect of prior knowledge on target-evoked P1 responses, suggesting that representations from prior knowledge activated by the cue interacted with target processing. Notably, the positive direction of this effect—increased pre-target alpha power predicted larger P1 amplitudes (Fig. 6)—directly contrasts with previous findings of a negative relationship between these variables ^68–70^, which is typically interpreted as reflecting the inhibitory nature of alpha rhythms ^71,72^. Indeed, our observation directly contrasts with the notion of alpha as a purely inhibitory or “idling” rhythm. We suggest that, in our task, increased pre-target alpha-band power may reflect the preactivation of neurons representing prior knowledge about object identity, thereby facilitating subsequent perceptual same/different judgments. This account is supported by the recent finding from invasive recordings in the Macaque that in inferior temporal cortex, stimulus-evoked gamma and multiunit activity are positively correlated with prestimulus alpha power, in contrast with the negative correlation observed in V2 and V4 ^73^.On the basis of this we speculate that the alpha modulation we observed in concert with P1 enhancement may have its origin in regions where alpha is not playing an inhibitory role.

Although our results are supportive of a general tenant of predictive processing accounts ^8,11,25^— that predictions, formed through prior knowledge, can influence sensory representations—our results also depart in an important way from certain proposals made by predictive coding theorists ^8,74,75^. With respect to the neural implementation of predictive coding, it is suggested that feedforward responses reflect the difference between the predicted information and the actual input. Predicted inputs should therefore result in a *reduced* feedforward response. Experimental evidence for this proposal, however, is controversial. Several fMRI experiments have observed reduced visual cortical responses to expected stimuli ^76–78^, whereas visual neurophysiology studies describe most feedback connections as excitatory input onto excitatory neurons in lower-level regions ^79–81^, which may underlie the reports of enhanced fMRI and electrophysiological responses to expected stimuli ^22,39,82^. A recent behavioral experiment designed to tease apart these alternatives found that predictive feedback increased perceived contrast—which is known to be monotonically related to activity in primary visual cortex— suggesting that prediction enhances sensory responses ^83^. Our finding that prior knowledge increased P1 amplitude also supports the notion that feedback processes enhance early evoked responses, although teasing apart the scenarios under which responses are enhanced or reduced by predictions remains an important challenge for future research.

## Additional Information

### Author contributions

J.S. collected and analyzed electrophysiological data, and wrote the manuscript. B.B. designed the experiments, collected the data, analyzed the behavioral data, and wrote the manuscript. B.R.P supervised electrophysiological data collection and analysis, G.L. conceptualized the experiments, analyzed the data, and wrote the manuscript.

